## Supplemental Figures for "Recurrent disruption of tumour suppressor genes in cancer by somatic mutations in cleavage and polyadenylation signals"

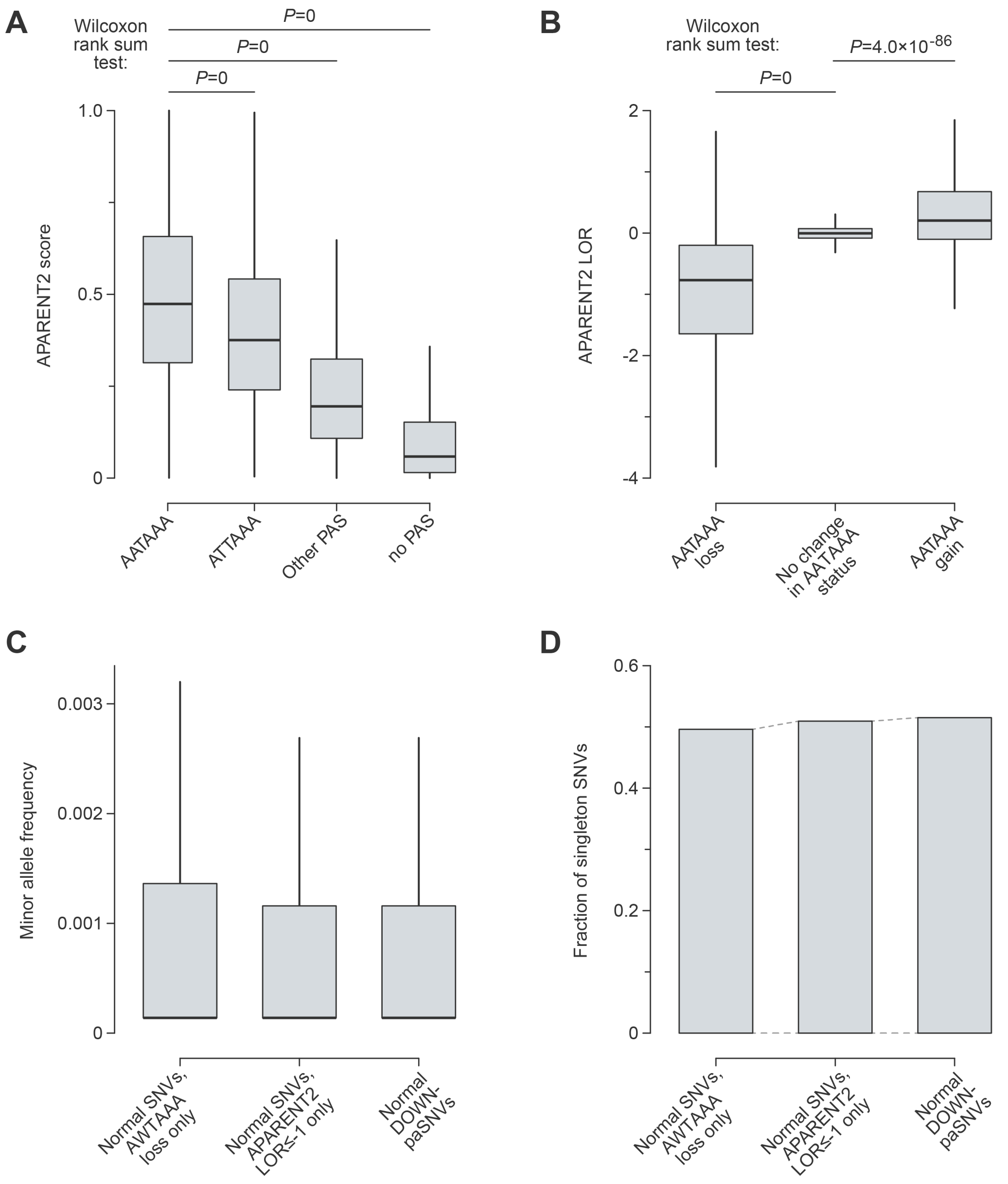


**Figure S1.** Distribution of cleavage/polyadenylation signal-disrupting mutations in the normal population (1000 genomes dataset).

**(A-B)** As expected, (A) AWTAAA-containing paSNVs tend to be associated with a relatively high APARENT2 score and (B) the loss or gain of AATAAA typically reduced or increased the score, respectively.

**(C)** Box plot comparison of normal-population allele frequencies of cleavage/polyadenylation signal-disrupting mutations defined by considering only AWTAAA gain/loss, only APARENT2 score changes, or both (DOWN-paSNVs).

**(D)** Bar plot comparison of normal-population fractions of singletons for cleavage/polyadenylation signal-disrupting mutations defined by considering only AWTAAA gain/loss, only APARENT2 score changes, or both (DOWN-paSNVs).

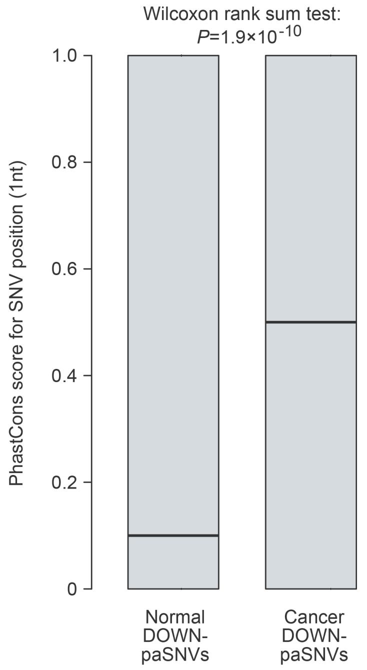


**Figure S2.** Cancer somatic DOWN-paSNVs often occur in evolutionarily conserved regions. The plot is generated similarly to Fig. 2D except the conservation was calculated for the exact SNV position.

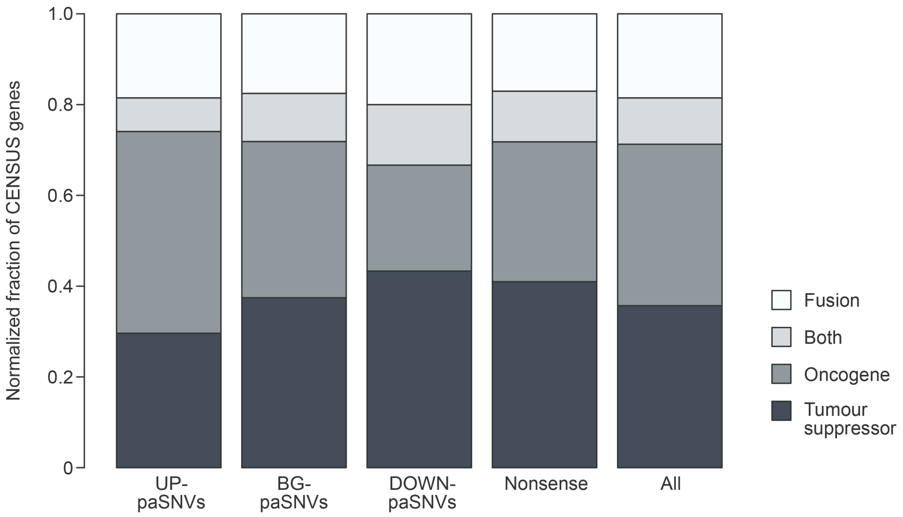


**Figure S3.** Cancer somatic DOWN-paSNVs often reside in genes with tumour suppressive functions.

Normalized stacked bar plot showing enrichment of SNVs disrupting polyadenylation signals (DOWN-paSNVs) in tumour suppressors for Census genes only. Note that nonsense mutations show a similar to DOWN-paSNVs enrichment in tumour suppressors, but not oncogenes. Conversely, UP-paSNVs are enriched in oncogenes but not tumour suppressors.

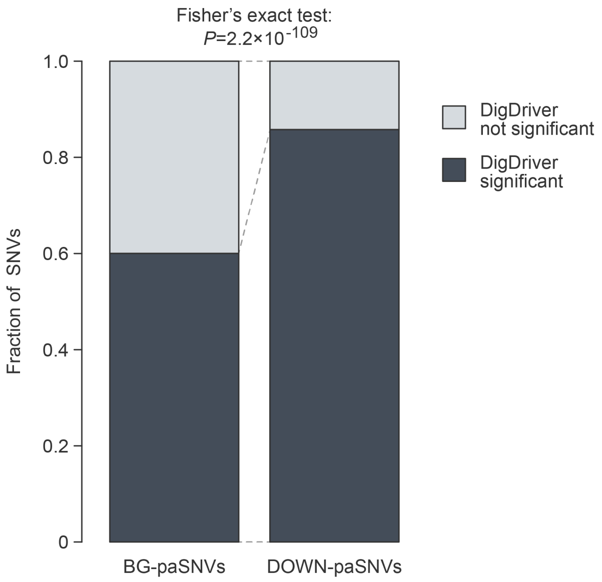


**Figure S4.** Cancer somatic DOWN-paSNVs are enriched for statistically significant DigDriver events (BH-adjusted $P<0.01$), suggesting that they may be under positive selection in cancer.

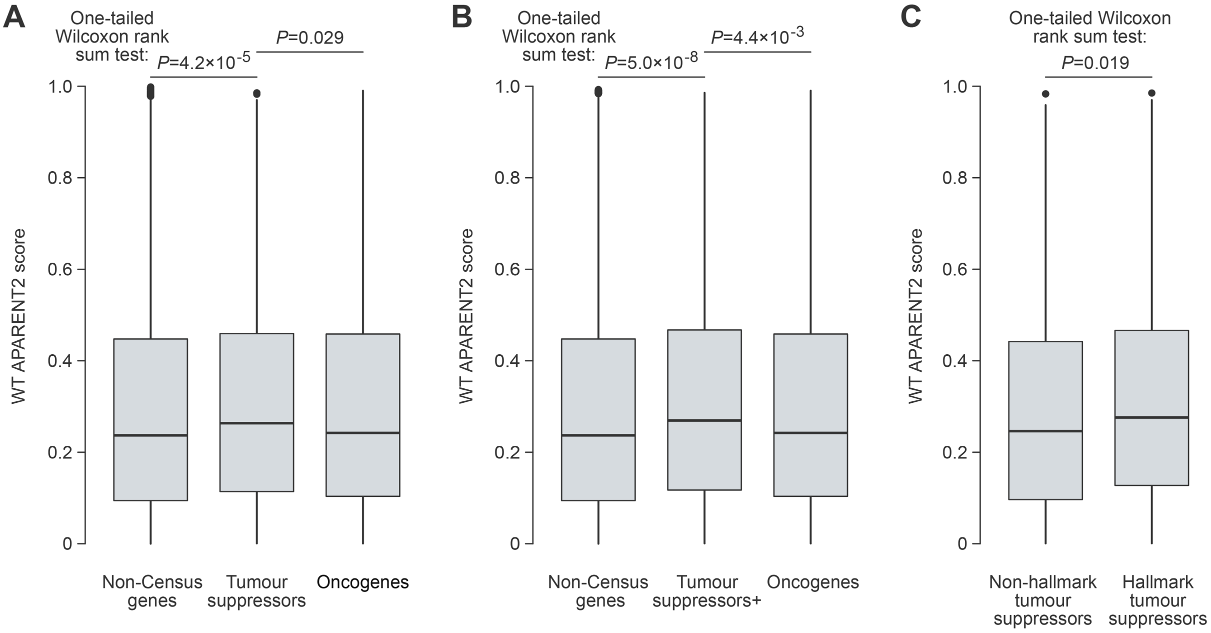


**Figure S5.** Wild-type tumour suppressor genes tend to have efficient cleavage/polyadenylation signals.

**(A)** Box plot showing that wild-type tumour suppressors have stronger cleavage/polyadenylation signals than oncogenes and non-Census genes.

**(B)** All Census genes classifiable as tumour suppressors ("Tumour suppressors+"; see Materials and Methods) have stronger cleavage/polyadenylation signals compared to oncogenes and non-Census genes.

**(C)** Tumour suppressors associated with "hallmarks of cancer" have stronger cleavage/polyadenylation signals than "non-hallmark" tumour suppressor genes.

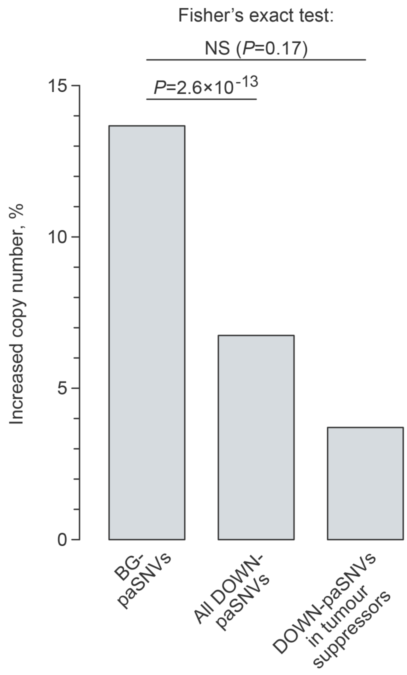


**Figure S6.** Reduced tendency for copy number increases in genes with DOWN-paSNVs.

The graph illustrates the fraction of genes with copy number increases >3 in at least one cancer sample. Notably, copy number increases are significantly less frequent in cancer genes affected by DOWN-paSNVs compared to their BG-paSNV counterparts. The lack of statistical significance in the comparison between tumour suppressors with DOWN-paSNVs and BG-paSNV genes is probably due to the limited number of CNV events in the former group.
